# Biomechanical properties of defence vibrations produced by bees

**DOI:** 10.1101/2024.01.15.575671

**Authors:** Mario Vallejo-Marin, David L. Field, Juan Fornoni, Daniel Montesinos, Cesar A. Dominguez, Ivan Hernandez, Gillian C. Vallejo, Charlie Woodrow, Ricardo Ayala Barajas, Noah Jafferis

## Abstract

**Abstract:** Bees use thoracic vibrations produced by their indirect flight muscles for powering wingbeats in flight, but also during mating, pollination, defence, and nest building. Previous work on non-flight vibrations has mostly focused on acoustic (airborne vibrations) and spectral properties (frequency domain). However, mechanical properties such as the vibration’s acceleration amplitude are important in some behaviours, e.g., during buzz pollination, where higher amplitude vibrations remove more pollen from flowers. Bee vibrations have been studied in only a handful of species and we know very little about how they vary among species. Here, we conduct the largest survey to date of the biomechanical properties of non-flight bee buzzes. We focus on defence buzzes as they can be induced experimentally and provide a common currency to compare among taxa. We analysed 15,000 buzzes produced by 306 individuals in 65 species and six families from Mexico, Scotland, and Australia. We found a strong association between body size and the acceleration amplitude of bee buzzes. Comparison of genera that buzz-pollinate and those that do not suggests that buzz-pollinating bees produce vibrations with higher acceleration amplitude. We found no relationship between bee size and the fundamental frequency of defence buzzes. Although our results suggest that body size is a major determinant of the amplitude of non-flight vibrations, we also observed considerable variation in vibration properties among bees of equivalent size and even within individuals. Both morphology and behaviour thus affect the biomechanical properties of non-flight buzzes.

**Summary statement:** Analyses across 65 bee taxa in three continents indicates that body size is a major determinant of the acceleration amplitude but not the oscillation frequency of non-flight thoracic vibrations.

## Introduction

Vibrations are key components of the behavioural repertoire of bees and have fundamental effects on bees’ survival and reproduction as well as on their ecological interactions with other organisms. Bees produce vibrations when powering wingbeat during flight, during intra-specific communication, to ward-off potential predators, and during pollen collection in certain types of flowers (Buchmann, 1985; Conrad and Ayasse, 2015; De Luca and Vallejo-Marin, 2013; Larsen et al., 1986). However, the biomechanical properties of these vibrations have been poorly studied beyond the context of flight (Dudley, 2000) and only in few bee taxa (e.g., Arroyo-Correa et al., 2019; Conrad and Ayasse, 2015; Hrncir et al., 2008; Jankauski et al., 2022; King et al., 1996; King and Lengoc, 1993). Given the enormous taxonomic diversity of bees (20,000 species, Almeida et al., 2023), and wide-ranging ecological, functional, morphological and size diversity (Danforth et al., 2019), establishing how variation in bee form permits and constrains the types of vibrations they produce is an important biological question.

Both flight- and non-flight vibrations are produced by rapid contraction of the indirect flight muscles that occupy most of the thorax capsule of bees and flies (Dickinson and Tu, 1997; Snodgrass, 1935; Snodgrass, 1942; Vallejo-Marin, 2022). In non-flight vibrations, although the thorax vibrates rapidly, the wings remain mostly undeployed. A type of thoracic vibration that appears to be common in bees and some flies, such as hoverflies (Syrphidae), is elicited when an individual is captured or manipulated, presumably as a mechanism to deter or startle potential predators (Moore and Hassall, 2016). This type of vibrations is often referred to as defence, annoyance, or alarm vibration (De Luca et al., 2014; Hrncir et al., 2008). In larger insects, defence vibrations have a clearly audible component and can be perceived as high pitch buzzes. In bumblebees, the entire nest engages in “hissing” when disturbed, and this aposematic signal can deter potential predators such as mice (Kirchner and Röschard, 1999). The biomechanical characteristics of non-flight vibrations have been little studied (e.g., Hrncir et al., 2005; Jankauski et al., 2022; King and Buchmann, 1996; King and Lengoc, 1993; Pritchard and Vallejo-Marin, 2020), but it is likely that these depend on both physiological and morphological characteristics of the insects that produce them. A hypothesis that can be used as a starting point, is that if muscle strain is scale-invariant, the displacement amplitude of defence vibrations should scale with thorax size. Furthermore, if the frequency of defence vibrations is relatively scale-invariant (which appears to be the case for pollination buzzes; De Luca et al., 2019; Vallejo-Marin, 2022), then we expect that both acceleration amplitude should increase with thorax size. Studies of wingbeat frequency show that, across species, there is a negative relationship between body size (e.g., mass) and wingbeat frequency (Dudley, 2000). In part, this relationship arises because wings with smaller areas need to be flapped at higher frequencies to generate enough lift compared to larger wings (Dickinson, 2006). In contrast, analyses of non-flight buzzes produced during buzz pollination (i.e., the pollination of flowers by bees using substrate-borne vibrations) suggest no relationship between bee size, measured as inter-tegular distance, and buzz frequency (De Luca et al., 2019; De Luca et al., 2014). It has been suggested that maximising buzz frequency (to the physiological maximum) during pollination might be advantageous because this would increase the vibration’s velocity amplitude for a given maximum displacement that the bee can achieve, and therefore dislodge pollen from anthers at higher rates (Vallejo-Marin, 2022). The functional consequences of buzz frequency on other non-flight vibrations in bees, such as defence buzzes, is unknown.

Bee size might also affect other properties of bee vibrations such as buzz duration. Empirical measurements of the duration of pollination buzzes in bumblebees suggests that larger individuals produce shorter buzzes (Barbosa et al., 2023; De Luca et al., 2014). However, this does not necessarily imply that larger bees are incapable of producing longer buzzes, and instead could simply reflect that a larger bee can extract more pollen with a short buzz than a smaller bee. In fact, larger bees may be able to buzz longer than smaller bees, as they have a lower mass-specific metabolic rate than smaller bees (Darveau et al., 2005), and longer buzzes may be advantageous when used in defence context. Thus, the relationship between bee size and buzz duration may be contingent on the behavioural context in which they are deployed.

In the present study, we conduct a survey of non-flight, defence vibrations by bees sampled in the field across three continents (the Americas, Europe, and Australia) to characterise the biomechanical properties of these vibrations in bees with diverse sizes and morphologies. Specifically, we address the question of how variation in thorax size affects the acceleration amplitude, frequency, and duration of defence buzzes. We hypothesise that: (1) assuming that thorax size reflects cross-sectional areas of indirect flight muscles (Clemente and Dick, 2023), acceleration amplitude of defence buzzes should scale positively with thorax size; (2) the duration of defence buzzes should be positively related to thorax size as larger bees have a lower-mass-specific metabolism and presumably can continue buzzing for longer than smaller bees all else being equal; and (3) to the extent that defence buzzes resemble pollination buzzes, any variation in the vibration frequency of defence buzzes is independent of thorax size.

## Materials and methods

### Bee sampling, identification, and estimate of thorax size

We conducted two new field expeditions to sample bee vibrations, one to western Mexico (Jalisco and Colima states; 20^th^ May – 5^th^ June 2022) and one to Australia (Western Australia and Queensland; 10^th^ September – 3^rd^ October 2022). In addition, we use an identical experimental procedure to re-analyse raw vibration data from bees that were sampled in Scotland (including the Outer Hebrides and Orkney) in May – August 2020. A subset of the Scottish vibration dataset has been previously published in a study comparing thoracic vibrations of bees and hoverflies (Vallejo-Marín and Vallejo, 2020) but here we re-analyse the whole bee dataset in order to maximise taxonomic and geographic coverage of our study. In each of these three geographic regions, we sampled bees visiting flowers, flying, or in surrounding vegetation using insect nets. Bees were placed individually in a plastic vial and stored in a cooler with ice packs. Bees were immediately transported to a field lab for measurement, and vibration data was acquired within 10 min to 3 h after collecting the bees. In Australia, we used a camper van as a mobile lab to minimise the time between sampling and data acquisition.

In Mexico and Australia, bees were photographed with an SLR camera and a macro lens (100mm f/2.8L, Canon, Tokyo, Japan), and the high-definition digital images used to determine thorax width size using ImageJ (Abramoff et al., 2004). In Scotland, the thorax width was measured with digital callipers (0.01 mm precision; CD-6-CSX, Mitutoyo Inc, Kanagawa, Japan). Thorax width was measured at the widest point of the thorax from a top view, usually at the height or just above the base of the forewing (tegulae). Linear measurements of the thorax, such as inter-tegular distance, are closely associated with other estimates of body size, including fresh and dry weight (Cane, 1987; Kendall et al., 2019) (Figure S1). For identification and sexing, voucher specimens were pinned and determined with the aid of entomological keys (G. Vallejo and M. Vallejo-Marin in Scotland; R. Ayala in Mexico; and K. Prendergast and T. Houston in Australia). Voucher specimens have been deposited in the National Museum of Scotland and the Queensland Museum, Australia.

### Vibration data acquisition

We used a miniature uniaxial piezoelectric accelerometer (0.2 g; 352C23, PCB Piezotronics, Hückelhoven, Germany) to measure vibrations produced by the bee’s thorax. The accelerometer was mounted at the end of a split bamboo stick (3.7 mm diameter x 200 mm length) with 30mm of connecting electrical cable (1 mm diameter, PCB Piezotronics) between the end of the stick and the accelerometer. The stick was affixed to a plastic box through two small holes. The natural resonance of the system was empirically measured to be approximately 17Hz (Vallejo-Marín and Vallejo, 2020). While the mass of the accelerometer is smaller than the largest bees measured, we note that it is much larger than that of the smallest bees. However, for defence buzzes, wing locking is expected to increase the bees’ resonance frequencies by approximately an order of magnitude compared with their flapping frequencies, so we expect the defence buzzes to be well below resonance, and thus in a regime dominated by the bees’ stiffness and damping, rather than inertia. In this regime, the added mass of the accelerometer would have only a small effect on the system dynamics. Future work will investigate this in more detail. To acquire the accelerometer signal we used a C-Series Sound and Vibration module (9250; NI, Newbury, UK) on a Compact DAQ chassis (cDAQ-9171, NI) connected to a portable computer. Accelerometer data was sampled at a rate of 10,240 Hz using custom software written in LabView NXG 5.0 (NI), stored as TDMS files (high throughput data format from NI) in the computer, and then converted to tab-separated text files for downstream analysis.

Bees were cold-anesthetised using icepacks or by briefly putting them in a -20°C freezer for 1-3 minutes until they became inactive. The bee was then tethered using a nylon loop (0.18 mm diameter) placed at the end of a blunt metal syringe needle (Pritchard and Vallejo-Marin, 2020; Vallejo-Marín and Vallejo, 2020). The bee was then allowed to return to room temperature and, once it had fully recovered, we pressed the dorsal surface of the thorax gently but firmly against the accelerometer so that the dorso-ventral axis of the thorax was aligned to the axis of vibration measurement of the accelerometer. We aimed to acquire vibrations for each bee during at least 45 s but the duration varied among individuals.

### Vibration analysis

Vibration data analyses were done in R ver. 4.3.1 (R Core Development Team, 2023) using the packages *seewave* (Sueur, 2018) and *tuneR* (Ligges et al., 2018). To analyse vibration data, we first applied high pass frequency filter (20 Hz, window length = 512) to remove background noise, and removed the offset by subtracting mean amplitude from each data file. We then used an automatic algorithm (*seewave:timer*) to identify all individual buzzes in a file with a detection threshold of 8% and a minimum duration of 100 ms. For some bees, the minimum duration was reduced to 50-60 ms to capture very short buzzes. Oscillograms and spectrograms of all the identified buzzes were examined by eye to remove spurious identifications caused by electrical noise or by accidentally knocking the accelerometer during bee manipulation. A full list of all buzzes identified, including those removed from downstream analysis, and their start and stop times relative to the start of the recording is provided in the supplemental data.

For each identified buzz, we calculated the fundamental frequency using the *seewave:fund* function with a window length of 512 samples, overlap of 50%, and a maximum frequency of 1000 Hz, and then obtained the median frequency for each buzz. For calculating vibration amplitude, we obtained the absolute amplitude envelope of the time series with a smoothing function window of two samples and overlap of 75% (*seewave:env*), and then calculated both peak and root mean square amplitude values for each buzz.

### Phylogeny and classification of buzz-pollinating genera

To incorporate phylogenetic relationships in our analysis, we used a genus-level phylogeny containing all the taxa studied here (Hedtke et al., 2013; tree 1). We chose a genus-level phylogeny as no equivalent species-level phylogeny exists for all studied species here, some of which are undescribed. Species were added to this phylogeny backbone as polytomies with branch length of zero. We also classified genera as either buzz pollinating or non-buzz pollinating using published information (Cardinal et al., 2018) and an unpublished list of buzz-pollinating taxa (A. Russell pers. comm.). A genus was considered to be capable of buzz pollination if at least one species in that genus has been reported to be buzz pollinating.

### Statistical analysis

We used linear mixed effects models using R ver. 4.3.1 (R Core Development Team, 2023) to investigate the relationship between vibration properties and bee size. We ran separate models for duration, fundamental frequency, and amplitude acceleration of buzzes. Thorax width was the explanatory variable and individual bee was used as a random effect. For buzz duration, we excluded buzzes shorter than 100ms (n=81 out of 150,000 buzzes) as this was the minimum buzz duration used for most individuals and prevents false positives when running the automated classification. The taxonomic and phylogenetic relationships among the taxa studied here were incorporated into the statistical analysis as follows. We used a Markov Chain Monte Carlo sampler for generalised linear mixed models that includes correlated random effects from a phylogeny, as implemented in the package *MCMCglmm* (Hadfield, 2010). In this model, we included species and individual as random effects, and phylogeny as the correlated random effects (pedigree). We ran 23,000 MCMC iterations with 3,000 discarded as burn-in and thinned every ten iterations.

## Results

We sampled 356 bees (81% females, 19% males), 202 in Scotland, 80 in Mexico and 74 in Australia. The 356 sampled bees belonged to 70 bee taxa in 6 families: Andrenidae (4% individuals), Apidae (74%), Colletidae (10%), Halicitdae (8%), Megachilidae (2%), and Stenotitridae (1%) (Figure 1). Of these sampled bees, 50 did not produce defence vibrations during our experiment (Table S1). Non-buzzing bees included *Tetragonula carbonaria* and *Trigona fulviventris,* in which no individual was observed buzzing (n > 8 individuals per species), and *Apis melifera* of which only a third of the sampled individuals produced a defence buzz (4/12). A full list of buzzing and non-buzzing individuals is provided in Table S1. In total, we extracted buzz data from 306 individual bees (80% females and 20% males), in 65 taxa from six bee families (Table S1). The thorax width of buzzing bees ranged from 0.84 to 9.6 mm (mean = 4.2 mm, SD = 1.6 mm). For bees in which both thorax width and inter-tegular distance was measured, the correlation coefficient was very high (Pearson’s rho = 0.985, *P* < 0.001) (Figure S1). Most buzzing bees were sampled in Scotland (60% of individuals) with the remaining nearly equally split in Mexico and Australia (21% vs 19%, respectively).

**Figure 1.**
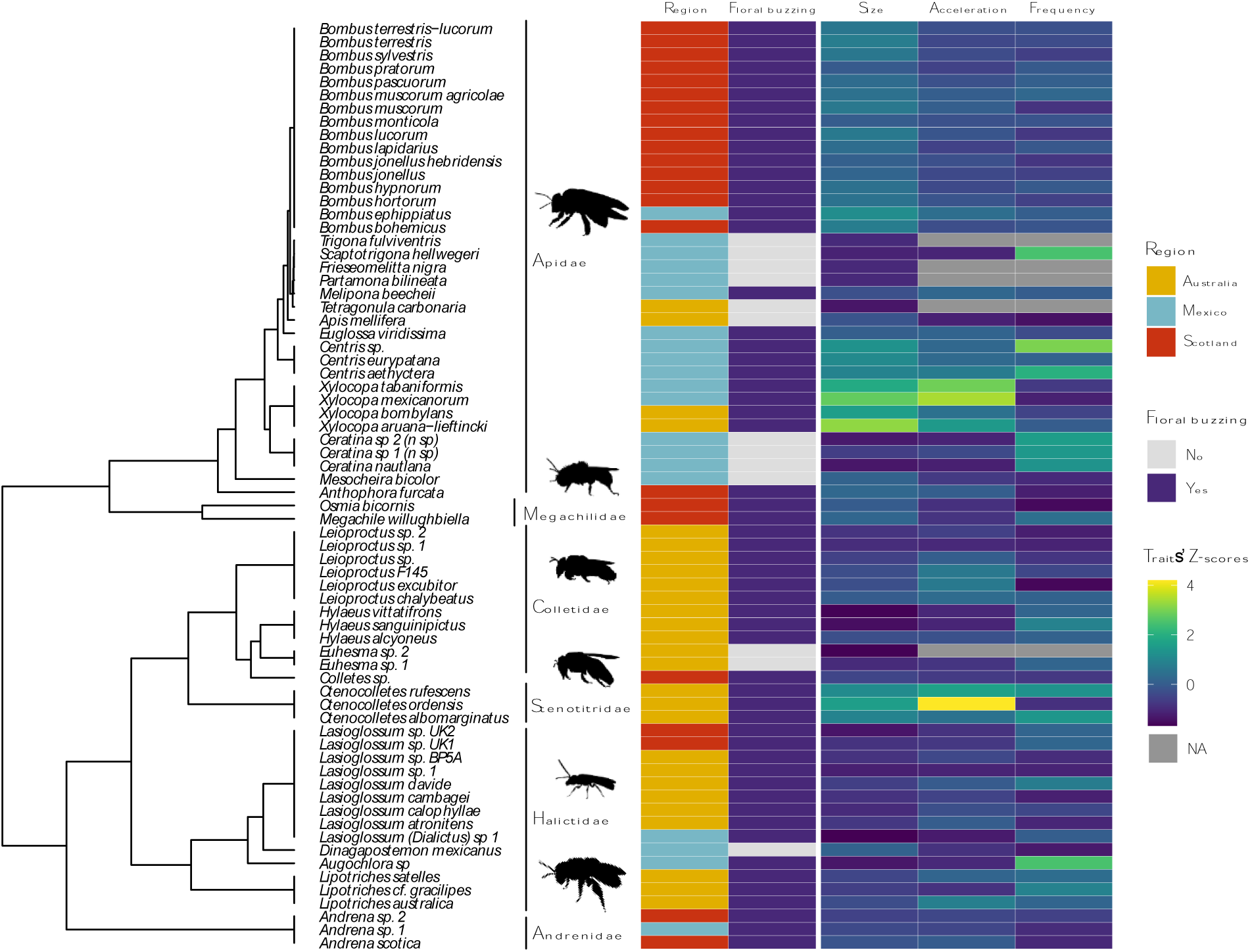
Phylogenetic relationships (at the genus level) among the bee taxa studied here showing variation in thorax width (size), vibration fundamental frequency (frequency), and peak acceleration per buzz. The first column indicates the sampled region, and the second column indicates whether the genus is known to buzz pollinate flowers or not. For ease of comparison, trait values (columns 3 to 5) are presented as Z-scores.

We recorded 15,000 individual buzzes, for an average of 49 buzzes per bee (median = 35 buzzes; range = 2 - 319). The average duration of a defence buzz was 0.76 s (range = 0.05 – 43 s; median = 0.29 s), and the average fundamental frequency was 205 Hz (range = 40 – 488 Hz; median = 193 Hz) (Figure S2). The peak acceleration amplitude of defence buzzes ranged from 2 – 1,473 m s^-2^, with an average of 112 m s^-2^ (approximately 11.4 g-force equivalents; median = 82.7 m s^-2^). Peak amplitude acceleration was strongly correlated with RMS amplitude acceleration (Pearson’s rho = 0.854, *P* < 0.001) (Figure S1).

We found that thorax width was positively associated with peak acceleration amplitude, as assessed in the MCMC model incorporating the phylogenetic structure of the data at the genus level and both species and individual identity as random effects (Figure 2, Table 1A). This model also indicated a statistically significant difference between buzz-pollinating and not-buzz pollinating taxa, with bees belonging to genera that are not known to buzz pollinate producing lower acceleration amplitude buzzes (Figure 2, Table 1A). In contrast, the fundamental frequency of non-flight buzzes was not associated with thorax width and did not differ between buzz pollinating and non-buzz pollinating taxa (Figure 2B, Table 1B). For buzz duration, we found a statistically significant effect of body size with larger bees producing longer buzzes (Figure 2, Table 1C). Whether a bee belongs to a buzz pollinating genus or not had no effect on buzz duration.

**Figure 2.**
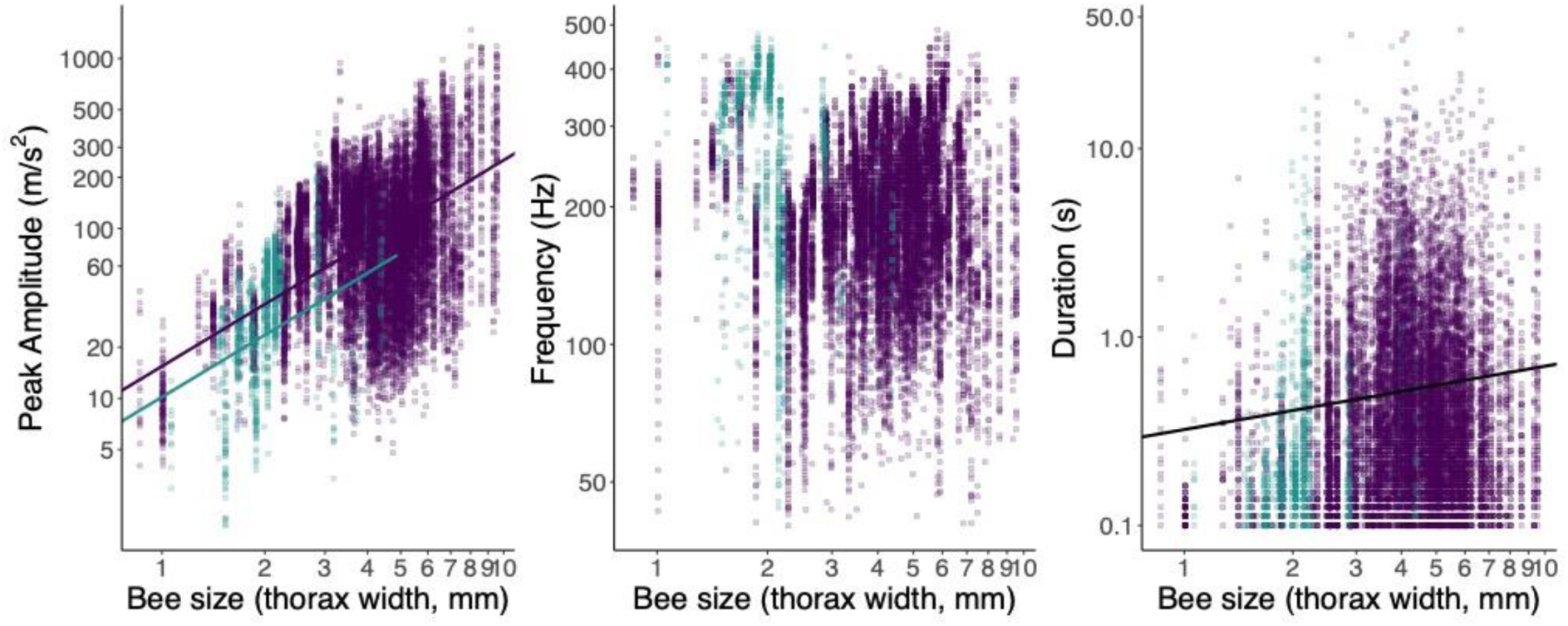
Bivariate plots of (A) peak amplitude acceleration, (B) fundamental frequency and (C) buzz duration against bee’s thorax width. The solid lines in panel A and C represent the regression models from Table 1. Each data point represents a single buzz. The colour of the symbols represents whether the data and regression line belong to a genus known to buzz pollinate (purple) or not (green).

**Table 1.** Model estimates of the of the effect of thorax width and whether a bee belongs to a buzz-pollinating genus or not on each of three response variables: peak amplitude acceleration (A), fundamental frequency (B), and buzz duration (C). Models were fitted with a Bayesian, MCMC approach using the *R package MCMCglmm* and with a genus-level phylogeny as correlated random effects and species and individual as random effects (see Methods: Statistical analyses). Ln = natural log.

| Response variable | Model coefficient | Estimate | 95% CI | $P_{MCMC}$ |
| --- | --- | --- | --- | --- |
| **A.** |  |  |  |  |
| **Ln(Peak amplitude acceleration)** |  |  |  |  |
|  | Intercept | 2.738 | 2.060 – 3.343 |  |
|  | Ln(Thorax width) | 1.213 | 1.004 – 1.402 | <.0001 |
|  | Non-buzz pollinating genus | -0.417 | -0.675 – -0.145 | <.0001 |
| **B.** |  |  |  |  |
| **Ln(Fundamental frequency)** |  |  |  |  |
|  | Intercept | 5.324 | 4.537 – 6.055 |  |
|  | Ln(Thorax width) | -0.112 | -0.294 – 0.058 | 0.222 |
|  | Non-buzz pollinating genus | 0.087 | -0.138 – 0.321 | 0.455 |
| **C.** |  |  |  |  |
| **Ln(Buzz duration)** |  |  |  |  |
|  | Intercept | -1.130 | -1.823 – -0.520 |  |
|  | Ln(Thorax width) | 0.336 | 0.077 – 0.591 | 0.013 |
|  | Non-buzz pollinating genus | -0.264 | -0.563 – 0.070 | 0.113 |

## Discussion

As expected, a main determinant of defence vibration amplitude is the size of the bee’s thorax, specifically the width of the mesosoma. A similar relationship between thorax size and vibration amplitude acceleration holds also for defence buzzes produced by hoverflies (Vallejo-Marín and Vallejo, 2020). This thorax width-amplitude relationship is probably related to an increase in the volume and cross-sectional area of the indirect flight muscles, but this hypothesis remains to be further explored. Thorax width should scale positively with the cross-sectional area of the dorso-ventral and dorsal longitudinal muscles (indirect flight muscles), which power bee buzzes across all behaviour contexts (Dickinson et al., 1998; Vallejo-Marin, 2022). In particular, if the proportion of muscles to body is scale-invariant, then the thorax width would be approximately proportional to the square root of the muscle cross-sectional area. We do not expect the characteristics of muscle tissue to vary too widely among different species of bees because they do not vary across asynchronous flight muscles of distantly related insect lineages (Marden 2000). Therefore, we conjecture that the positive relationship detected here between thorax width and vibration amplitude reflects the increased power of larger indirect flight muscles in larger bees. In particular, if the muscle contraction is limited by some scale-invariant strain, the muscle deformation amplitude would scale linearly with thorax width, and, given the observed scale-invariant buzzing frequency, this would result in the acceleration amplitude also scaling linearly with thorax width, which is very close the measured data. Additionally, if the muscle properties, proportion of muscles to body size, and body shape are scale-invariant, then we would expect muscle stiffness to scale linearly with thorax width, and the muscle force to scale with the square of thorax width, which would also result in a linear scaling of amplitude with thorax width, for operation in a stiffness-dominated regime.

While these hypotheses are consistent with our measured data, they remain to be further explored. It is unknown whether higher amplitude defence vibrations are more effective at deterring potential bee predators such as birds or mammals. However, vibration amplitude directly affects the amount of pollen removed from flowers during buzz pollination. Our results suggest that, all else being equal, a larger bee should be able to remove more pollen per time spent buzzing than a smaller bee. This does not mean that smaller bees are at a disadvantage in terms of total pollen removal, as the behaviour of the bee on the flower will affect how these vibrations are applied to the flower and the amount of pollen removed, and small bees might adjust how they manipulate and interact with flowers to maximise pollen removal (Barbosa et al., 2023; Delgado et al., 2023).

Bees belonging to buzz pollinated genera produced higher amplitude vibrations than those not in buzz-pollinated genera. If a bee is unable to produce vibrations of sufficient amplitude to dislodge pollen, this could explain why some bees do not buzz pollinate (e.g., *Apis mellifera*)(King and Buchmann, 2003). However, the vibration amplitude of defence buzzes in buzz pollinating and non-buzz pollinating bees shows considerable overlap, and it is unclear whether the statistical difference detected here has any functional effect on the amount of pollen removed from flowers. Indeed, this biomechanical limitation hypothesis does not seem to be a general explanation of why some insects do not buzz pollinate, as hoverflies (Diptera: Syrphidae), which are not generally known to buzz pollinate, are capable of producing vibration amplitudes equivalent to similarly sized buzz pollinating bees (Vallejo-Marín and Vallejo, 2020). Additional work is thus required to establish whether limits on the vibration amplitude that a bee can reach explains why some bees do not buzz pollinate.

The frequency of non-flight defence buzzes was not related to bee size. This is consistent with previous analyses of pollination buzzes (another type of non-flight thoracic vibration). Acoustic surveys of pollination buzzes among North American (De Luca et al., 2019) and neotropical bees (Burkart et al., 2011) found no statistical association between intertegular distance and buzz frequency, in contrast to the frequency of flight buzzes which scales negatively with size. In contrast to flight, defensive and floral buzzing allows the mechanics of the thoracic muscles to be uncoupled from both the aerodynamic properties and the size of the wings (Hickey et al., 2022), removing the frequency limitations enforced during flight. Frequency variation between buzzes produced by the same individual can also be mediated by bee behaviour. Bees may be able to alter the resonant properties of the thorax by using axillary muscles and altering thorax stiffness, as in the case of dipteran indirect synchronous muscles that regulate wing stroke frequency and amplitude (Dickinson et al., 1998). In addition, it is possible that the thorax could be driven at frequencies different to their resonant frequency, and there is some evidence that this is the case in the alarm and communication buzzes produced by *Melipona seminigra* (Hrncir et al., 2008), providing an additional mechanism to dynamically alter the frequency of thoracic vibrations (Vallejo-Marin, 2022). Together with previous work, our results suggest that across a wide diversity of bees, the frequency of non-flight buzzes, including pollination and defence buzzes is not strongly constrained by bee size (Burkart et al., 2011; De Luca et al., 2019). Therefore, bees appear to be relatively free to exploit different buzz frequencies regardless of their size, although we still do not fully know the functional consequences of frequency variation in the different behavioural contexts.

We observed a large amount of variation in the properties of buzzes of bees of similar size, and even within individuals. Similarly to vibration frequency, vibration amplitude is also partially under neural control of the bee, and the bee might have the capacity not only of turning the indirect flight muscles on and off, but also to modulate their amplitude similarly to the modulation of wing stroke amplitude in flies (Dickinson et al., 1998). The ability to control both amplitude and frequency of thoracic vibrations should help the bee to regulate energy expenditure, as the thoracic power muscles are capable of producing significant power (>100 W kg^-1^; Josephson, 1997) but are very energetically demanding. These potential mechanisms of regulating the timing, frequency, and amplitude of thoracic vibrations as well as the observed variation in buzz properties within individuals and between individuals and species of similar size, suggests that there is considerable flexibility in any potential constraints of morphology on vibration production. This flexibility in producing vibrations of different mechanical properties should be ecologically advantageous, as thoracic vibrations are deployed in very different contexts from flight to pollination to defence to mating (Conrad and Ayasse, 2015; King and Buchmann, 1996; Kirchner and Röschard, 1999; Larsen et al., 1986; Russell et al., 2017).

Despite the behavioural control of vibration properties, ultimately the limits of these properties are set by biomechanical and physiological constraints. The next step in understanding the capacity of a bee to produce non-flight buzzes should be to compare the architecture of the indirect flight muscles across species. Does cross-sectional area of the muscle scale isometrically with the width of the mesosoma? Do buzz pollinating taxa have larger muscles? How do different castes differ in their indirect flight muscle anatomies? Such questions offer a rich avenue for interdisciplinary research in bee phylogenetics, form, function, and biomechanics.

## Acknowledgements

We thank M. Saunders, M. Hall, M. Whitehead, F. Montealegre, and G. Martin-Ordas for their support in the development of the project and for their contribution in securing funding for fieldwork. Estación Biológica de Chamela-UNAM and Estacion Cientifica Las Joyas kindly provided access and support during fieldwork. We thank José Rubén Pérez Ishiwara for logistic support, and K. Prendergast and T. Houston for their assistance in identifying Australian bees.

## Data availability

Raw vibration data and the script fort statistical analyses will be deposited in a public repository (e.g., Dryad) upon acceptance.

## Supplemental Figures

**Figure S1.**
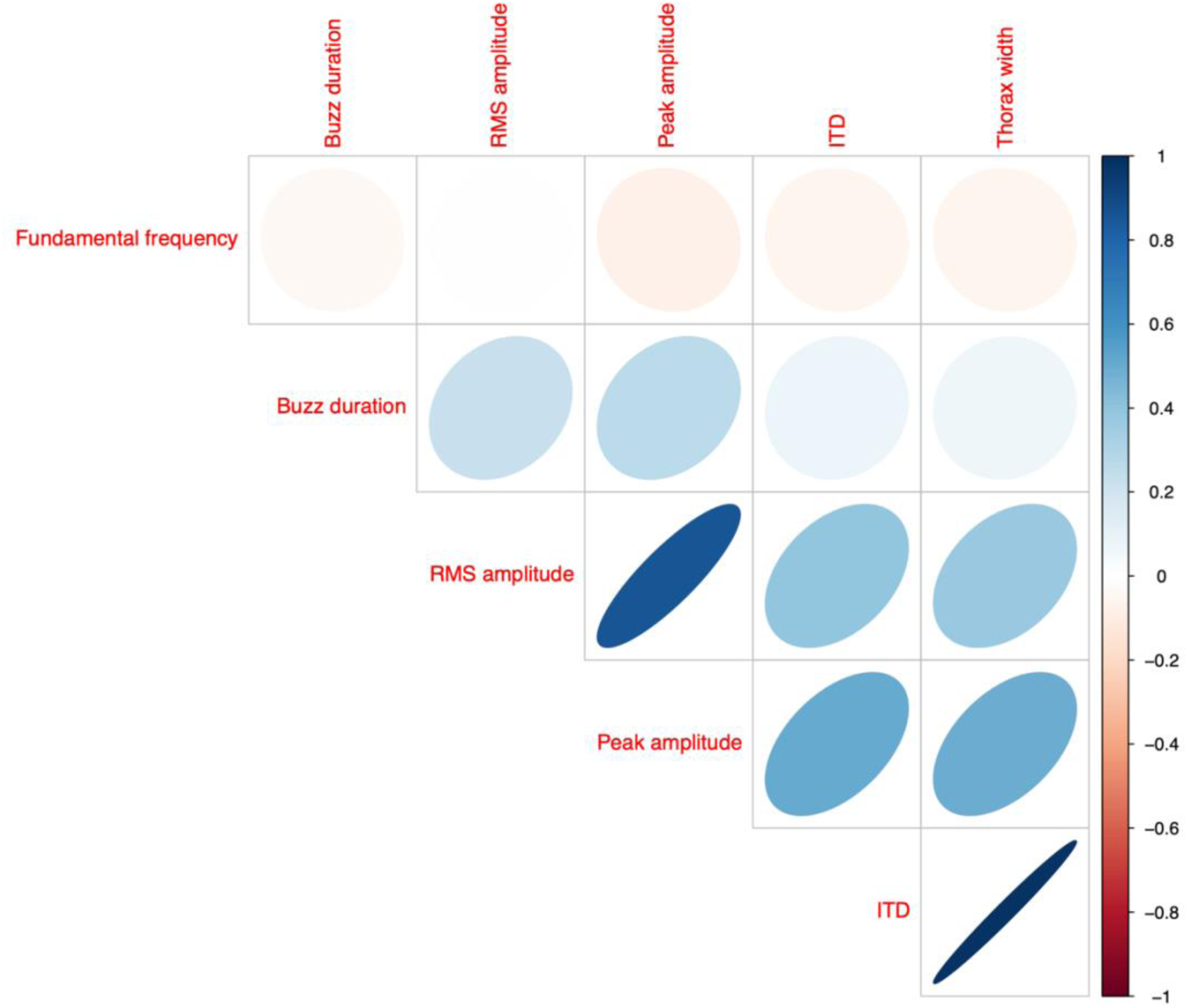
Correlation plot among buzz duration (s), root mean squared amplitude (RMS amplitude, m s^-2^), peak amplitude (m s^-2^), intertegular distance (mm), and thorax width.

**Figure S2.**
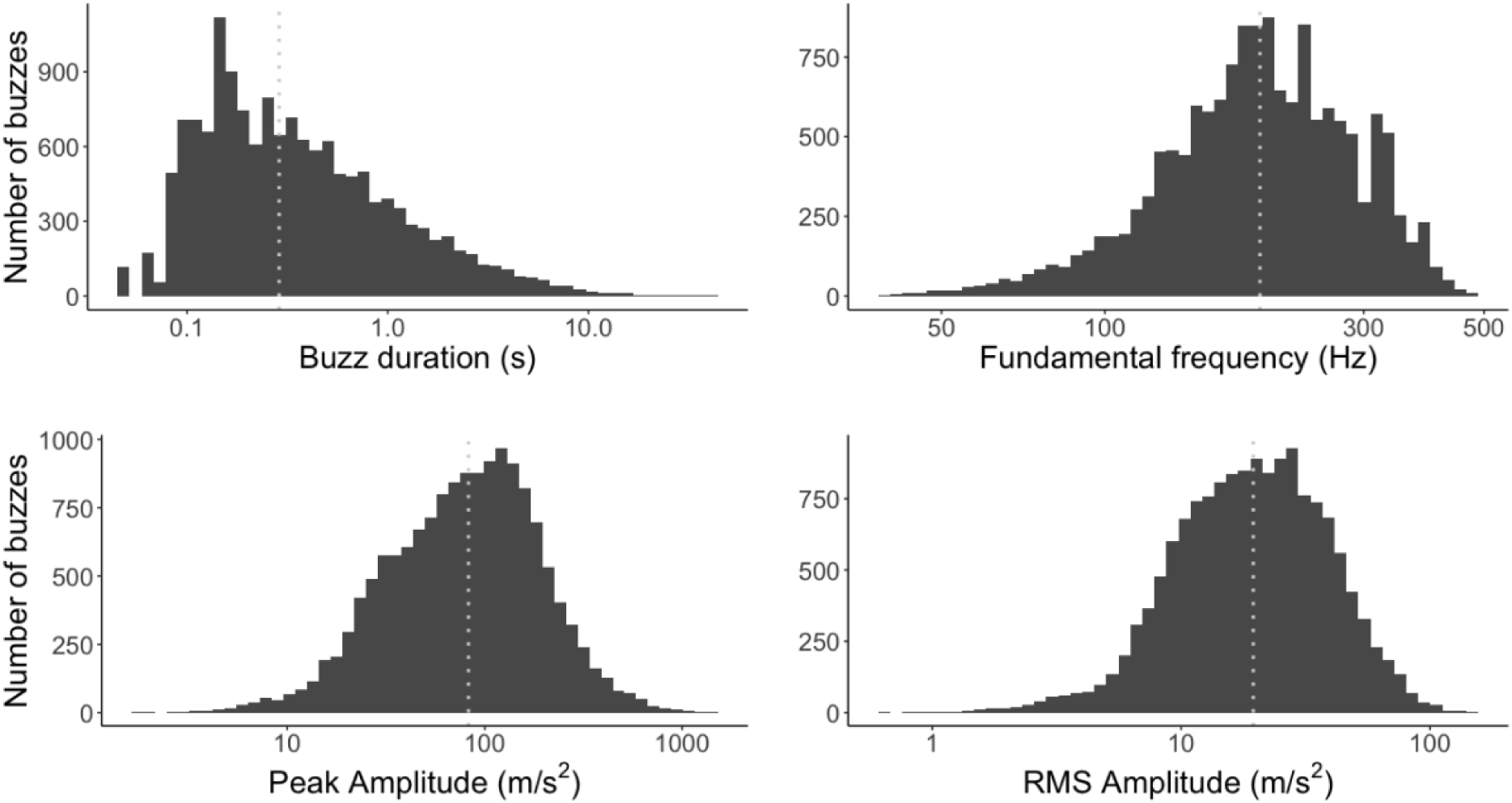
Histograms of the duration, fundamental frequency, peak amplitude acceleration, and root mean squared acceleration (RMS) of defence buzzes. X-axis is in log-10 scale. The vertical dotted lines represent median values. Median buzz duration = 0.288 s; median fundamental frequency = 193 Hz; median peak amplitude = 82.7 m s^-2^; RMS amplitude = 19.5 m s^-2^.

**Table S1.**
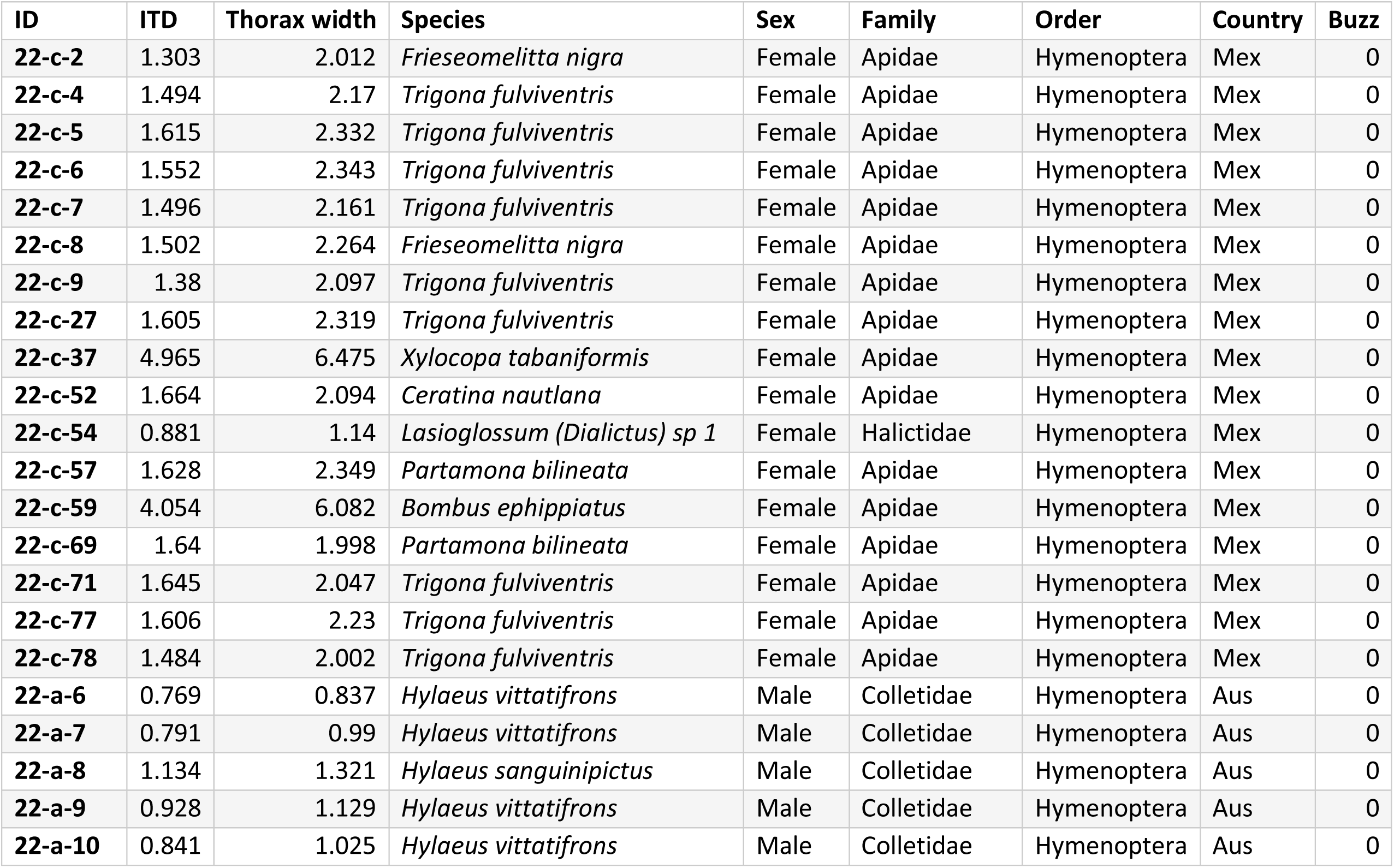
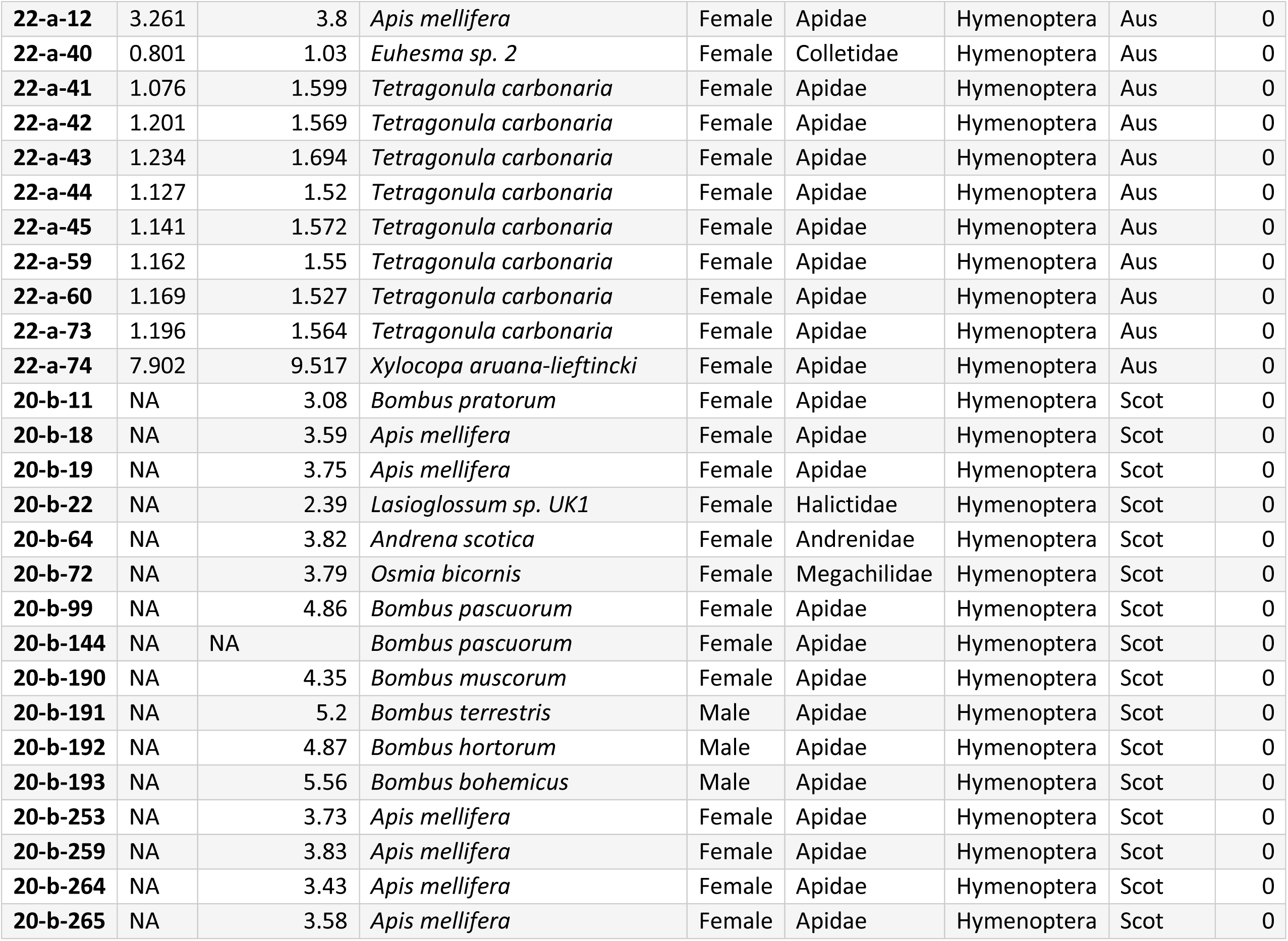
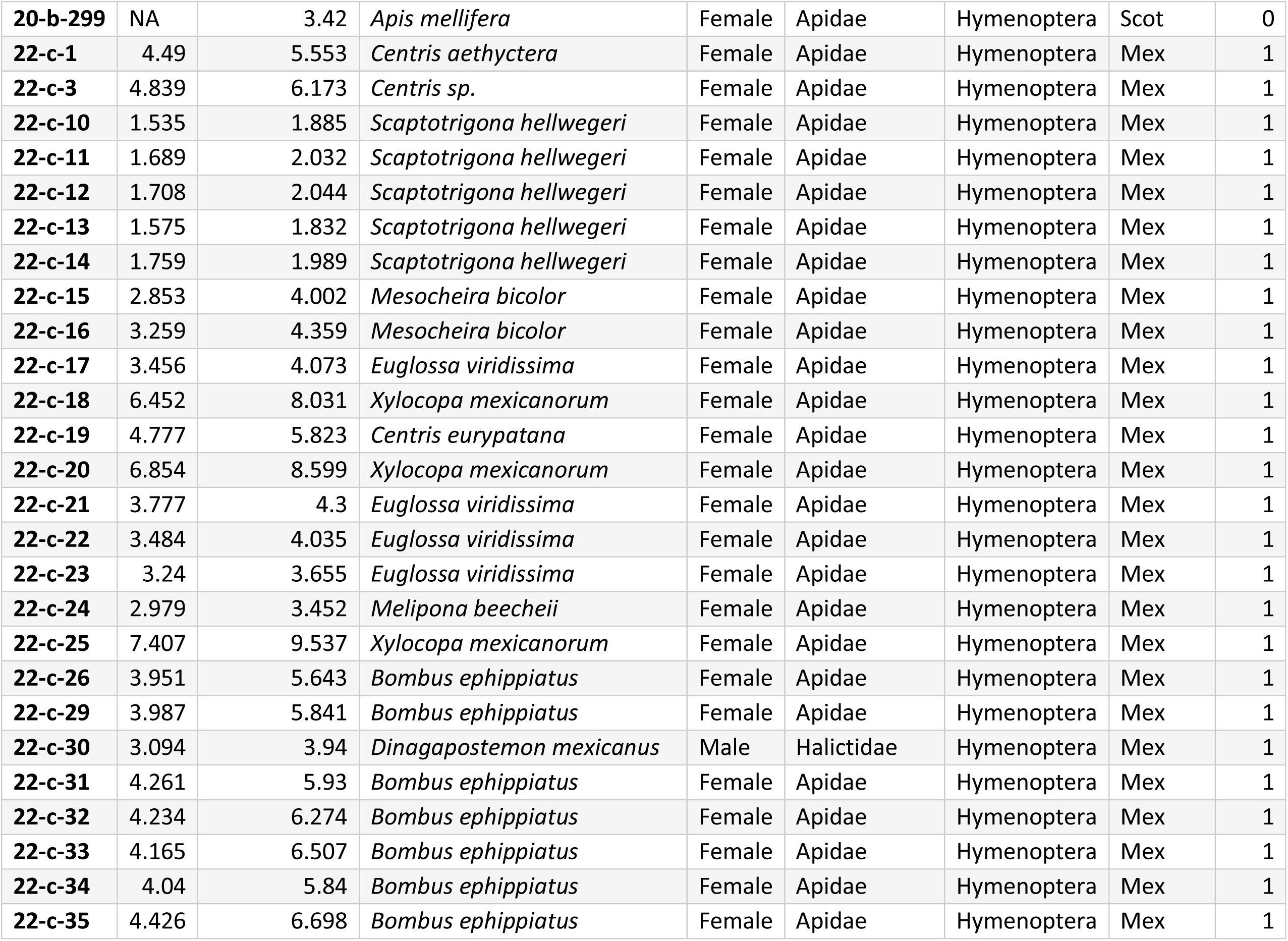
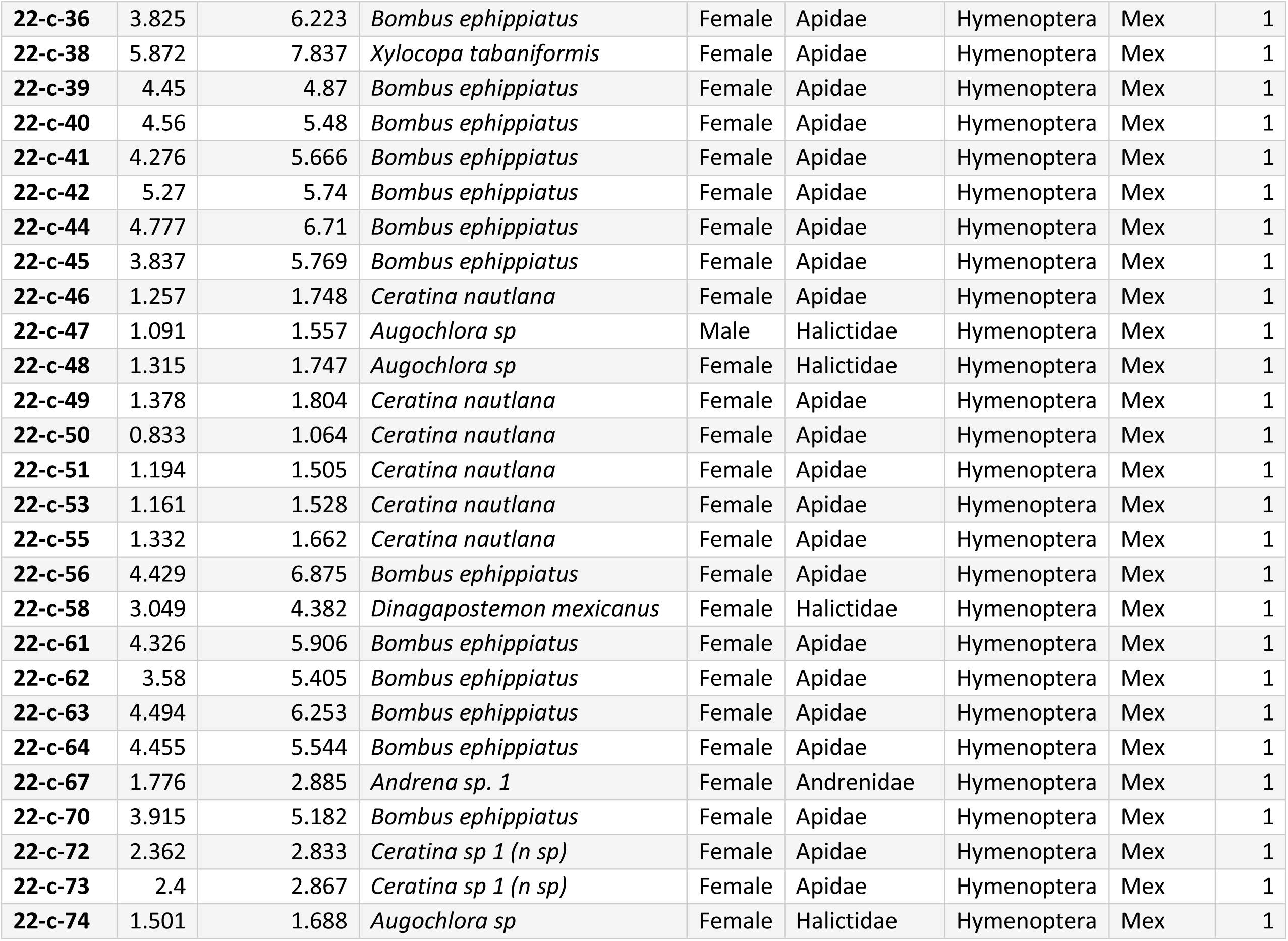
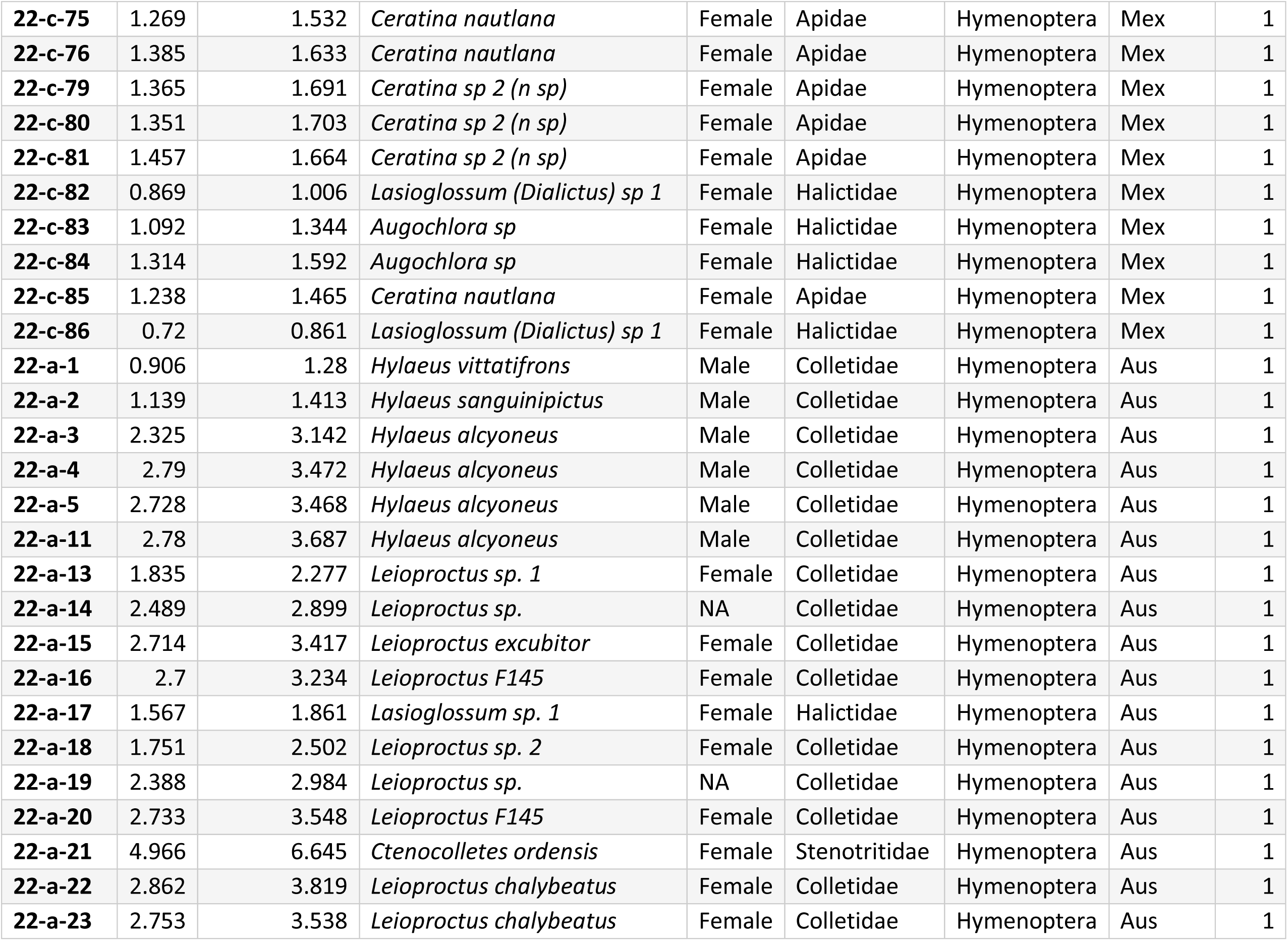
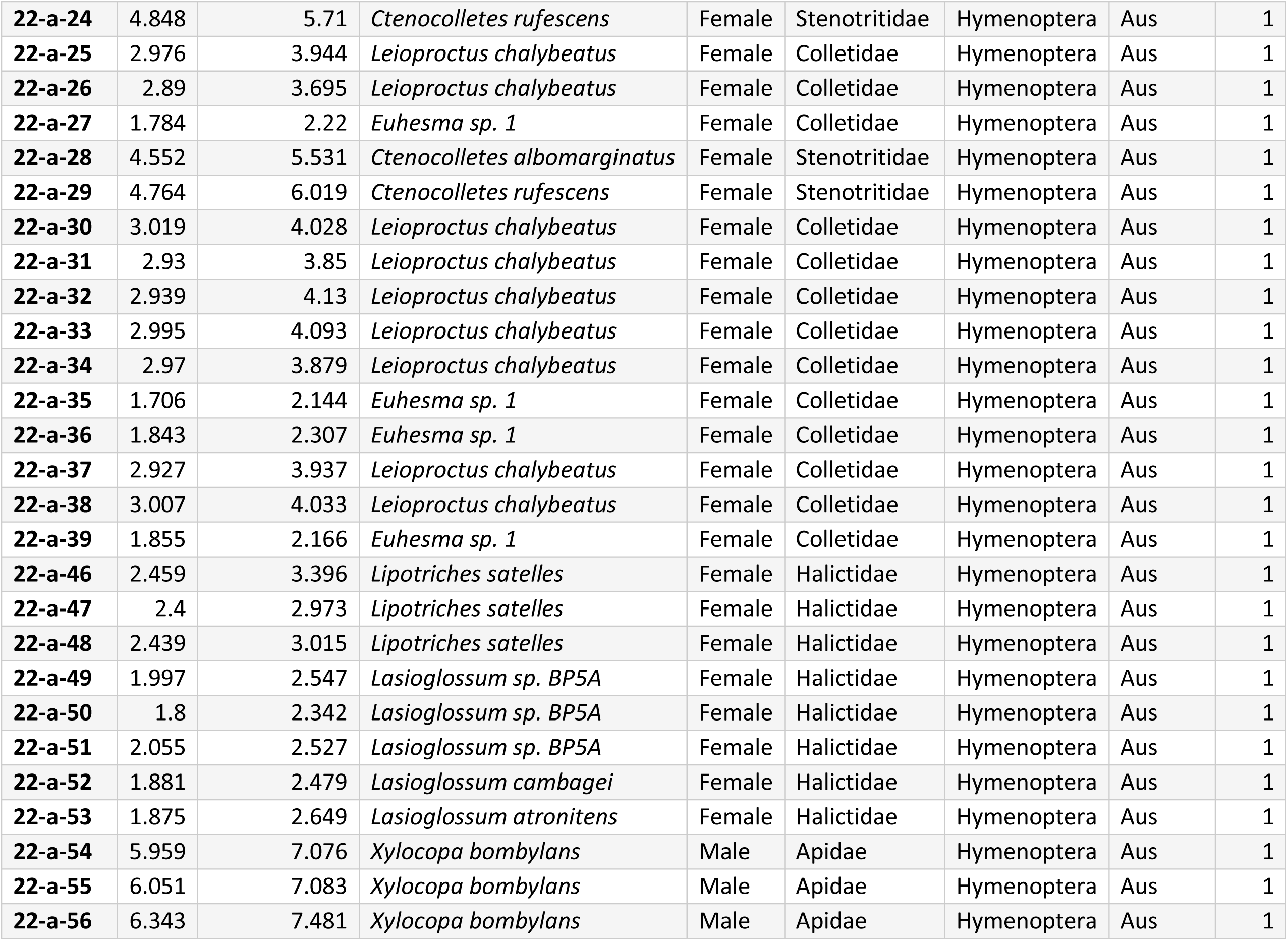
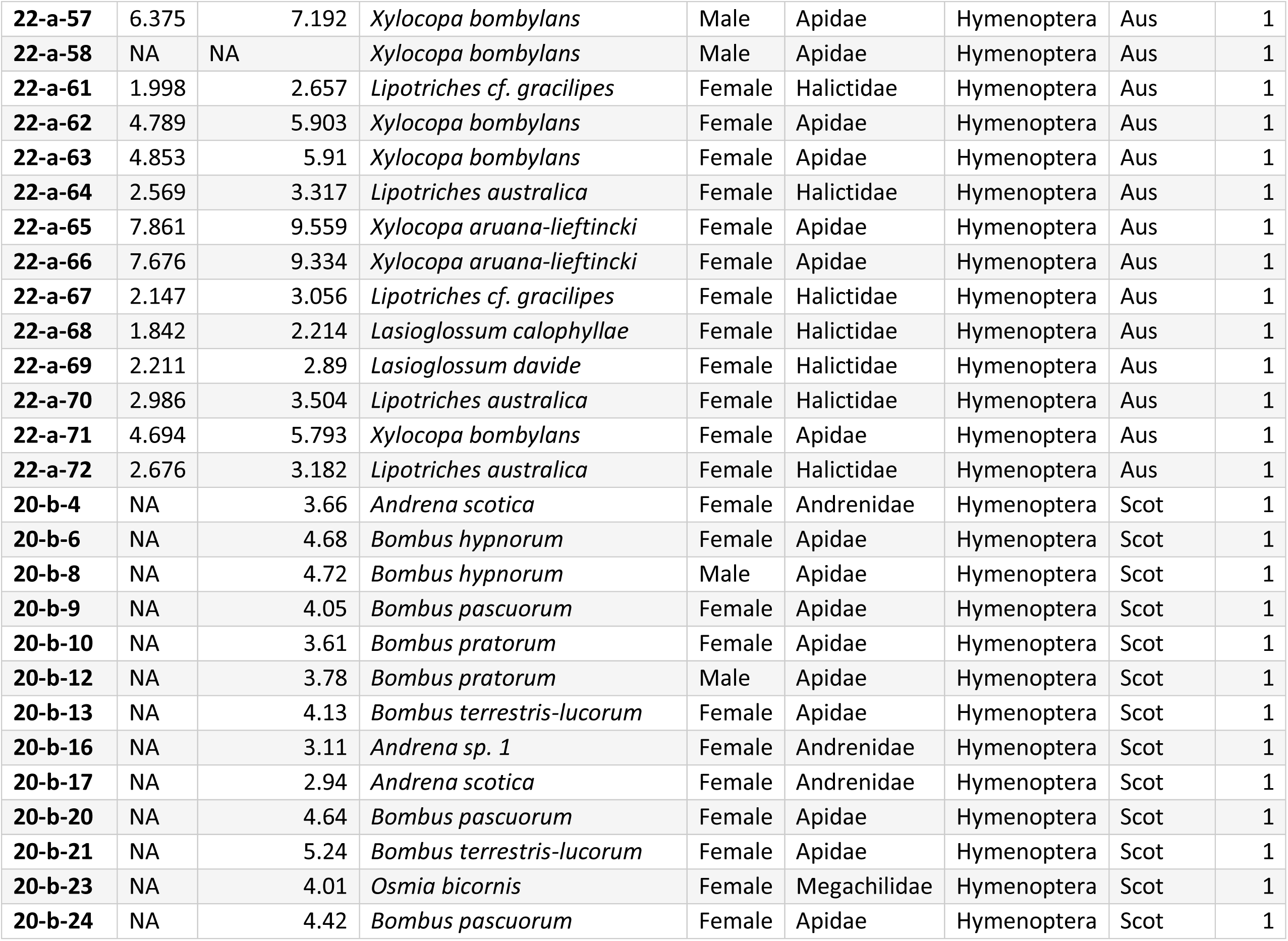
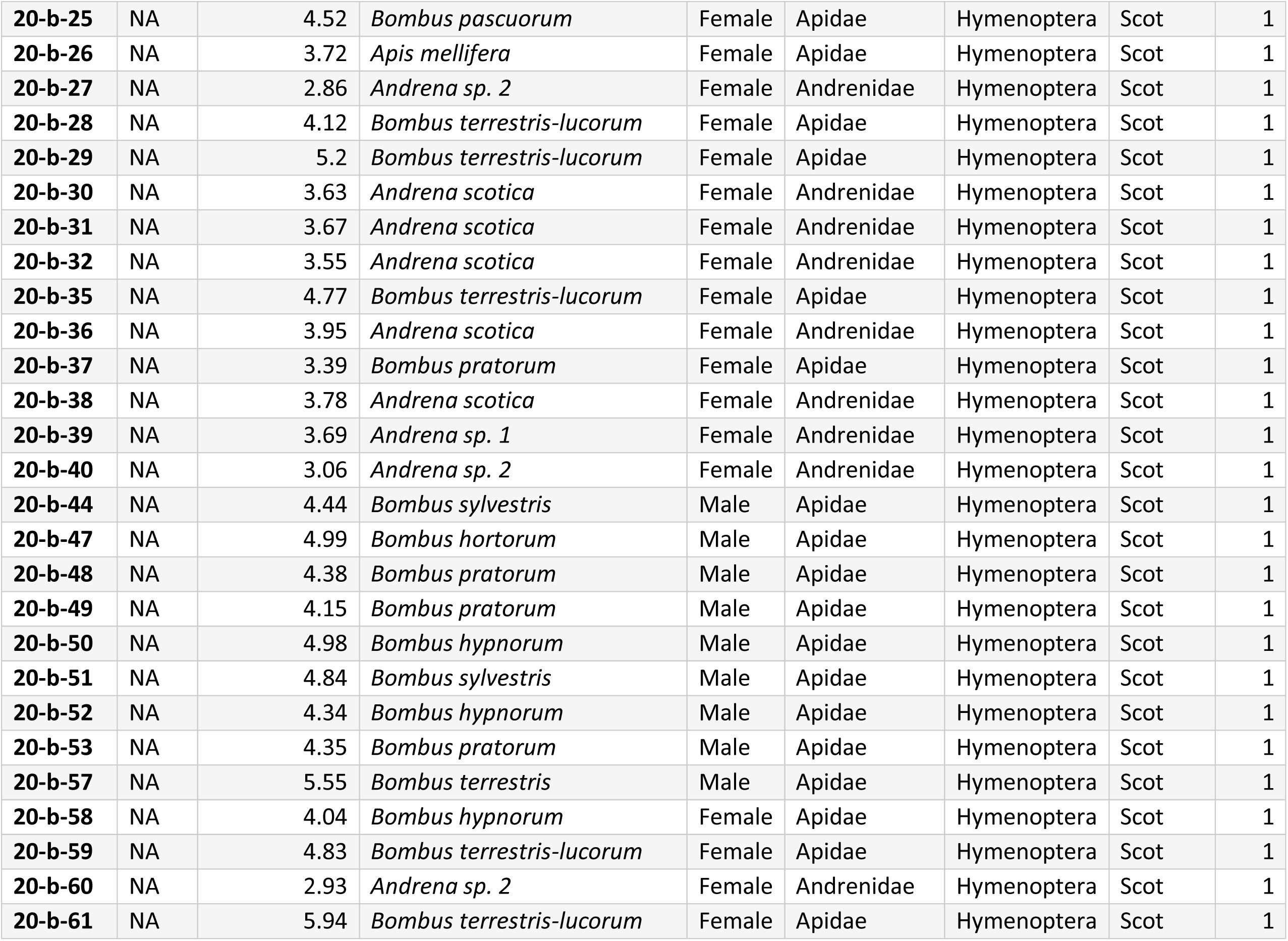
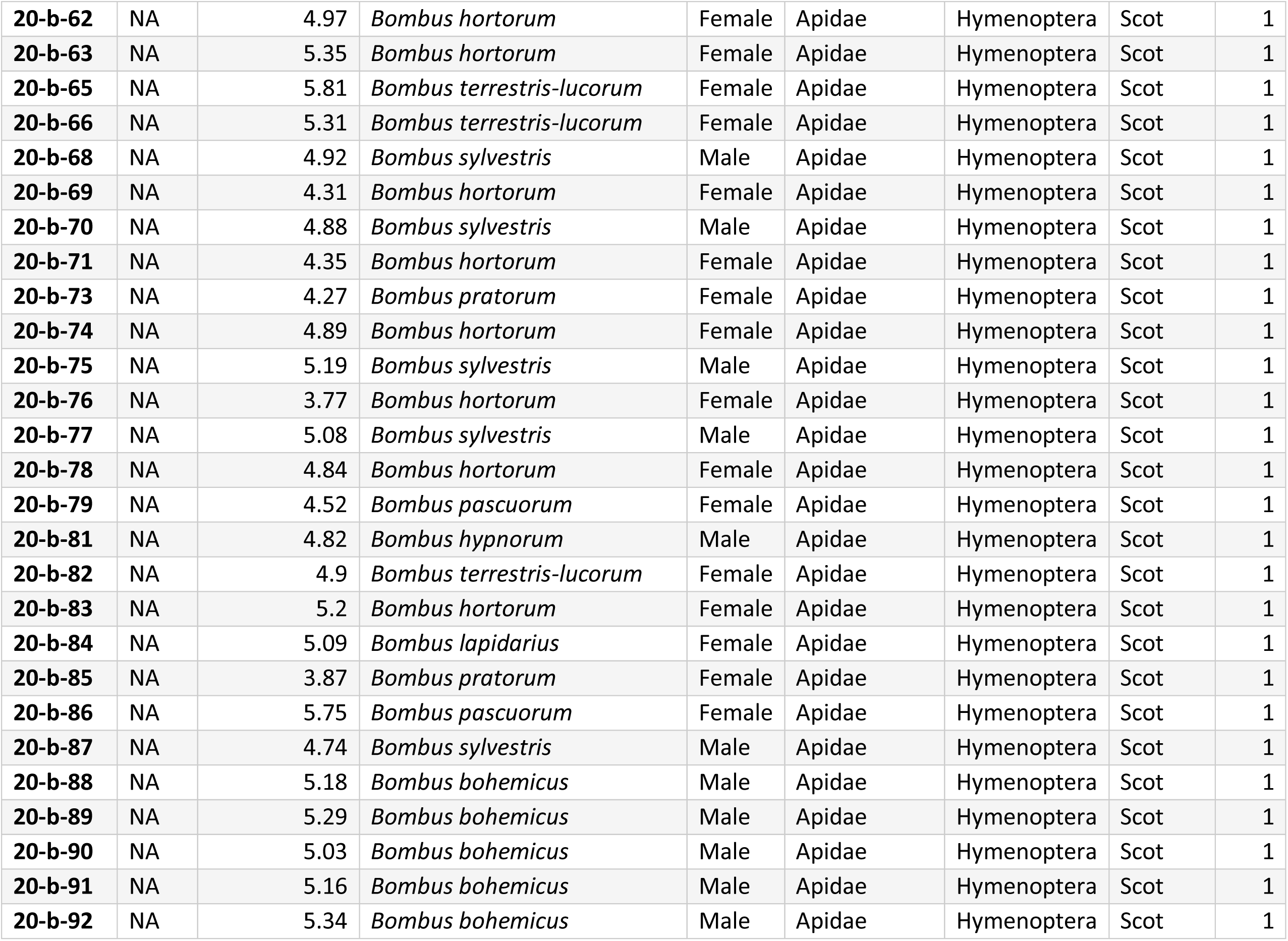
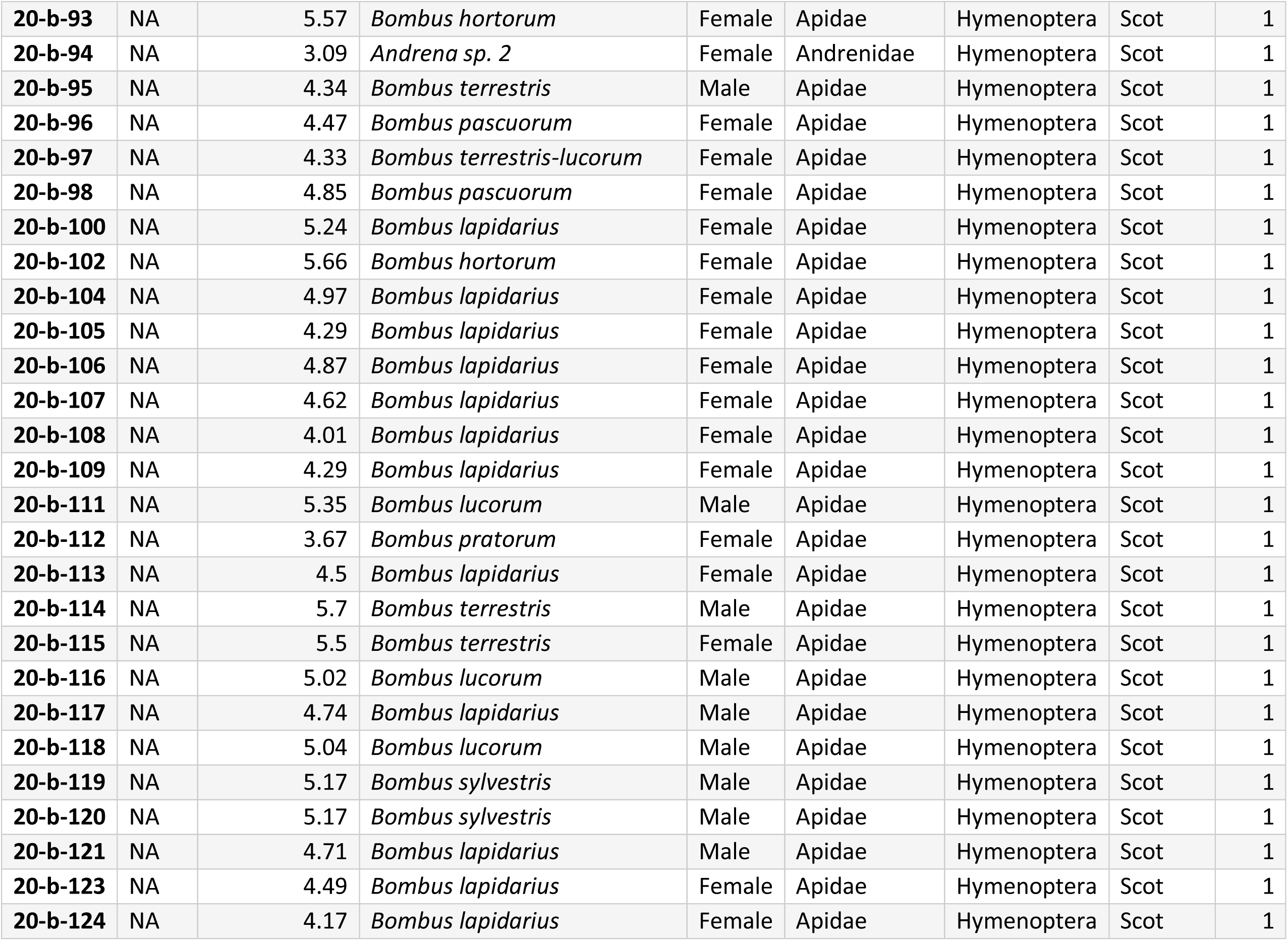
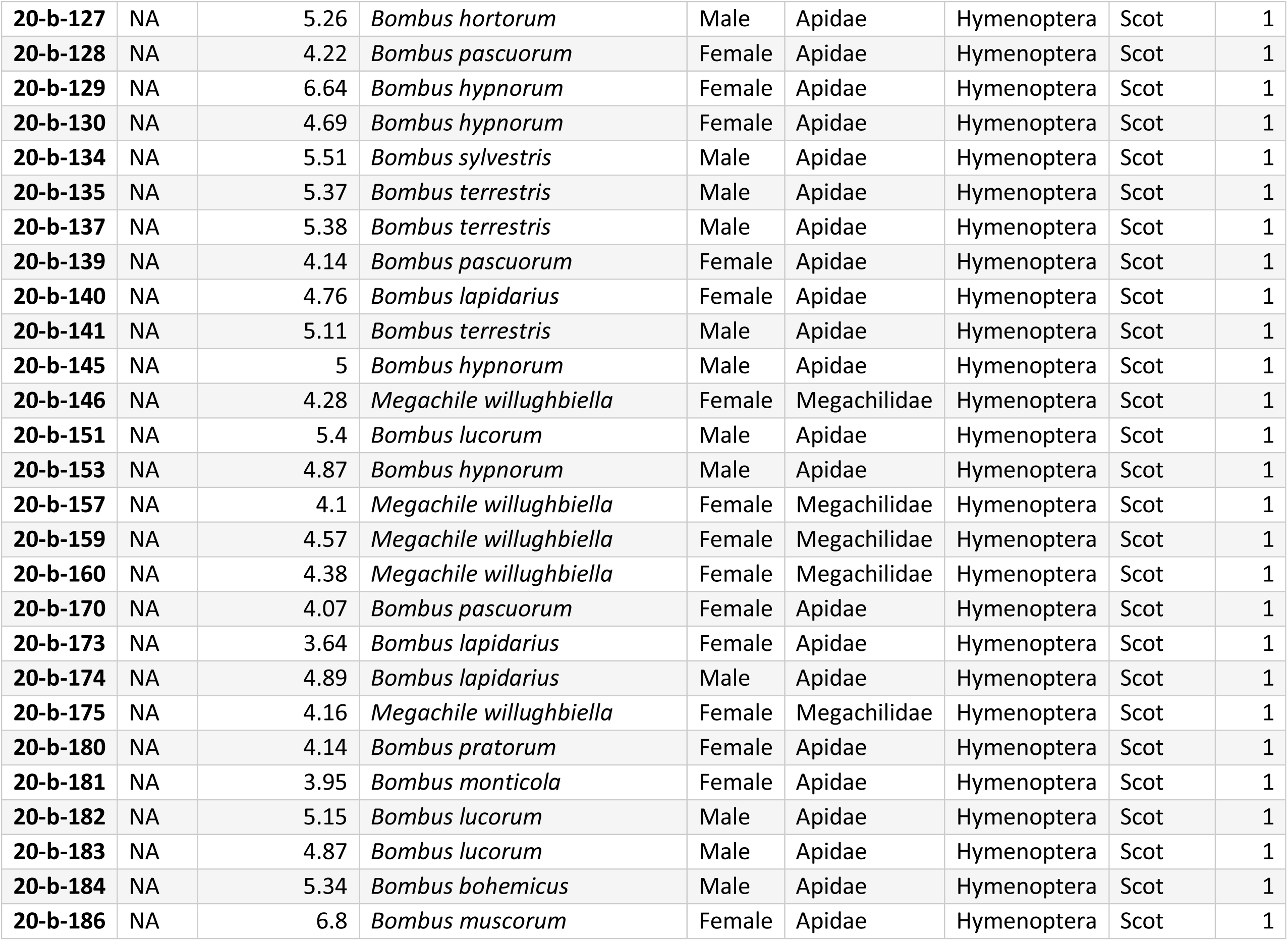
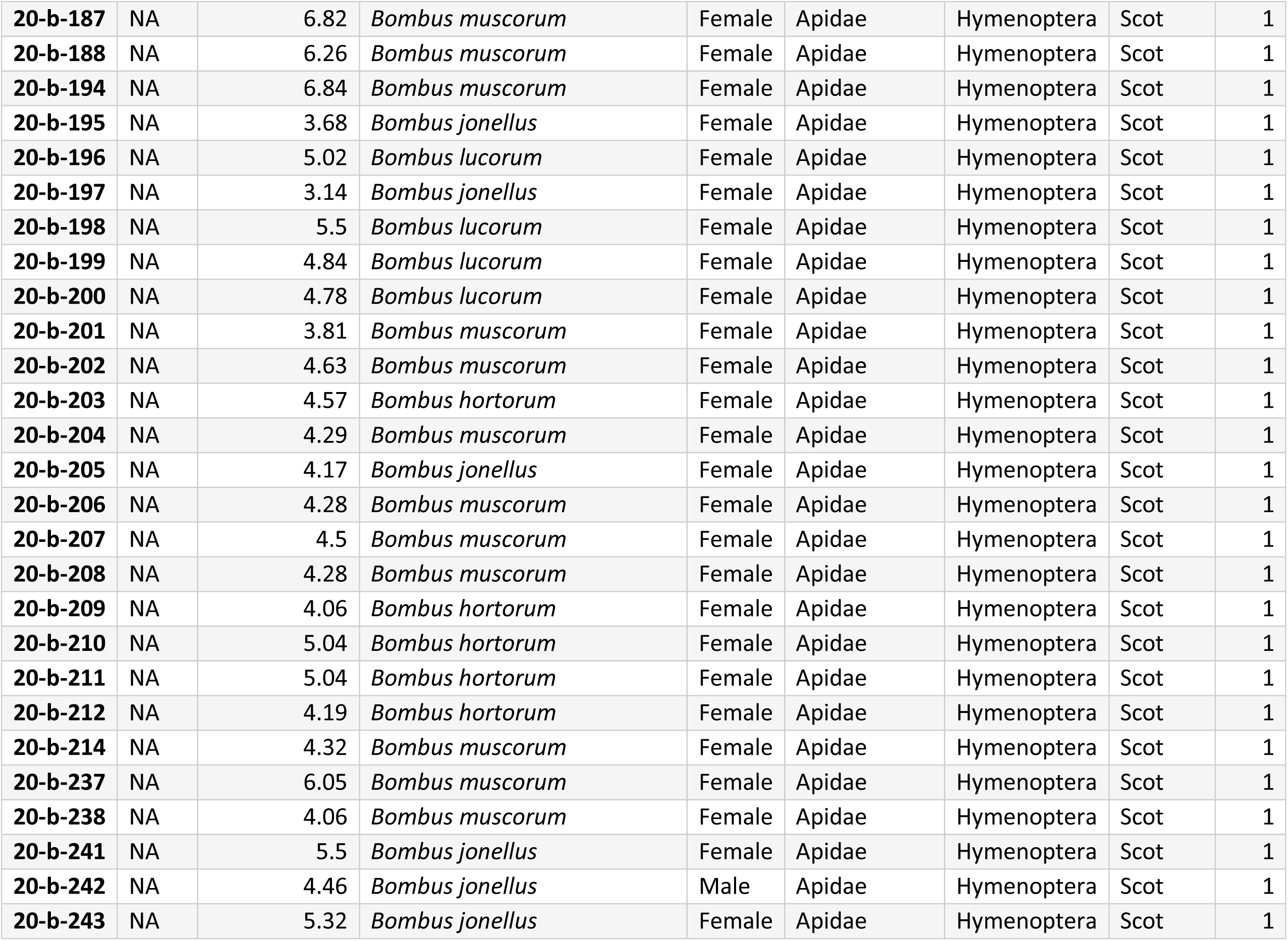
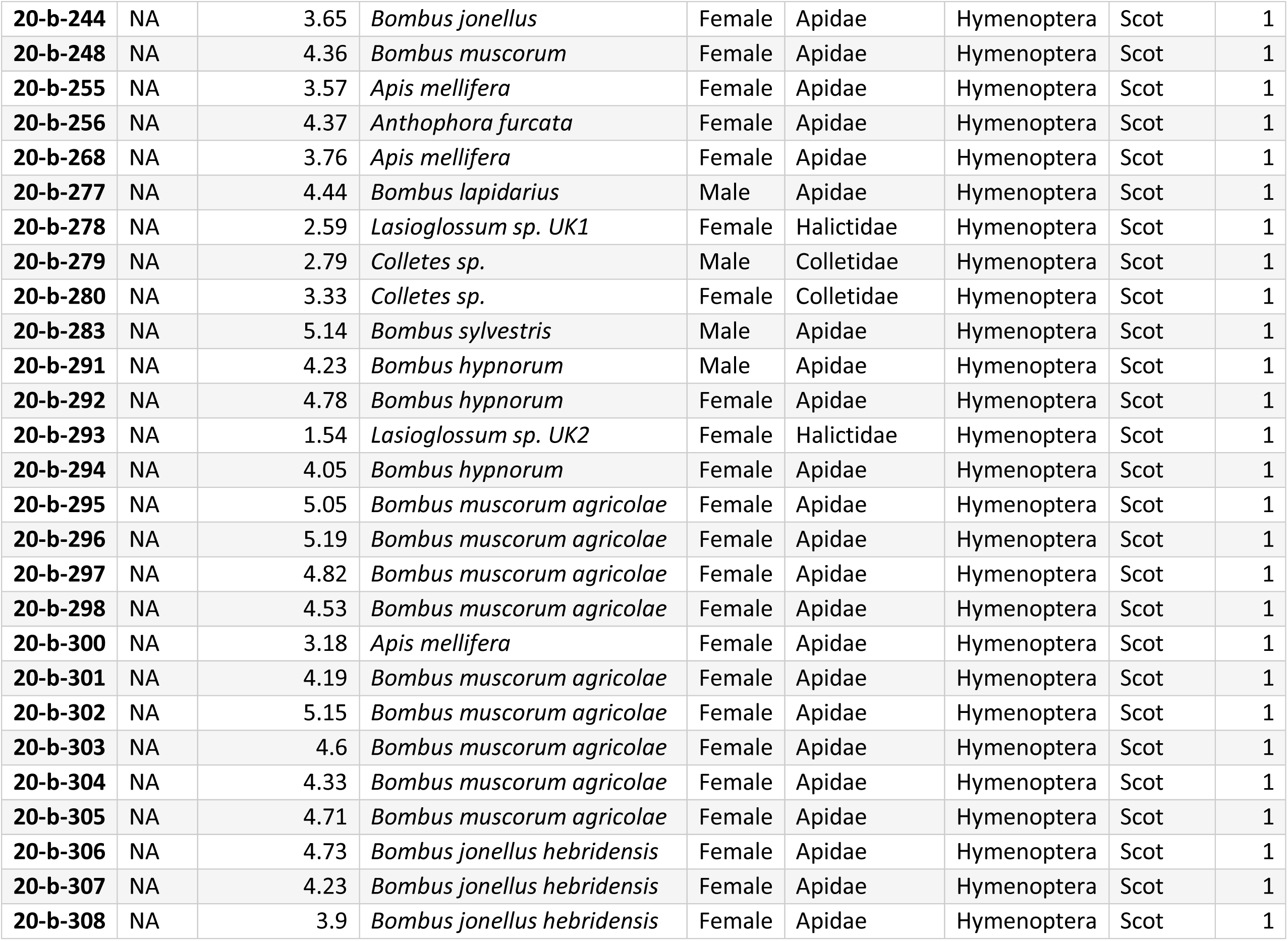
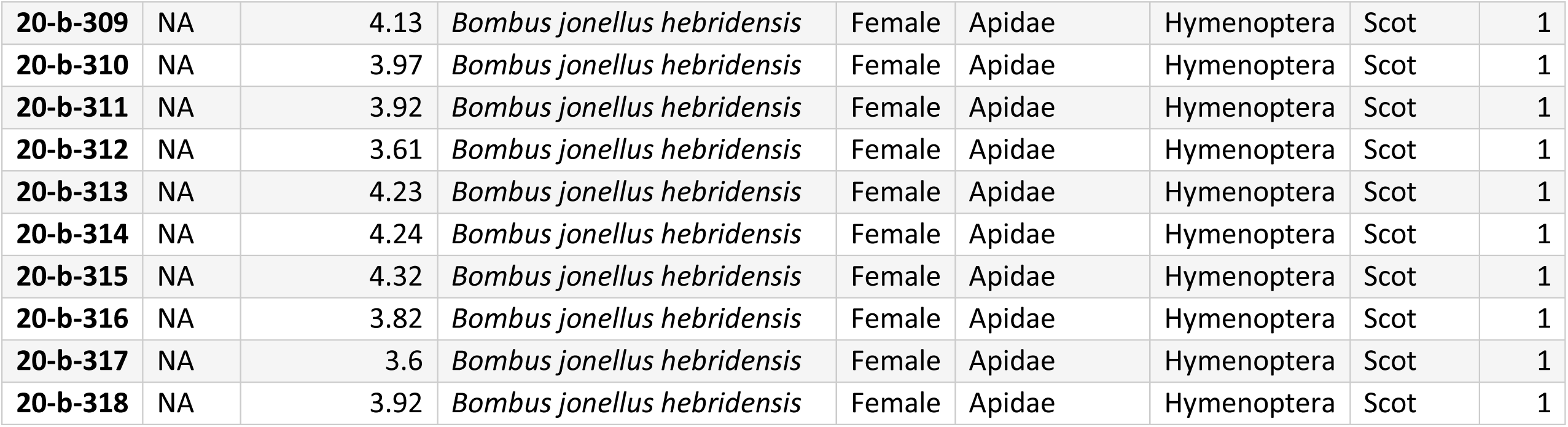
List of bees sampled. ID = Bee identification code, ITD = Intertegular distance in mm, Thorax width = thorax width in mm, Country = country sampled, Buzz = whether the bee produced a defence buzz during the experiment (1) or not (0).

## References

1. Abramoff, M. D., Magelhaes, P. J. and Ram, S. J. (2004). Image processing with ImageJ. Biophotonics International 11, 36–42.

2. Almeida, E. A., Bossert, S., Danforth, B. N., Porto, D. S., Freitas, F. V., Davis, C. C., Murray, E. A., Blaimer, B. B., Spasojevic, T. and Ströher, P. R. (2023). The evolutionary history of bees in time and space. Current Biology.

3. Arroyo-Correa, B., Beattie, C. and Vallejo-Marin, M. (2019). Bee and floral traits affect the characteristics of the vibrations experienced by flowers during buzz pollination. Journal of Experimental Biology 222, jeb198176.

4. Barbosa, B. C., Delgado de Lima, T. D., Mota, G. V. and Nogueira, A. (2023). The role of intraspecific variation in bumblebee body size and behavior on buzz pollination of a tropical legume species. American Journal of Botany, e16236.

5. Buchmann, S. L. (1985). Bees use vibration to aid pollen collection from non-poricidal flowers. Journal of the Kansas Entomological Society, 517–525.

6. Burkart, A., Lunau, K. and Schlindwein, C. (2011). Comparative bioacoustical studies on flight and buzzing of neotropical bees. Journal of Pollination Ecology 6, 118–224.

7. Cane, J. H. (1987). Estimation of bee size using intertegular span (Apoidea). Journal of the Kansas Entomological Society 60, 145–147.

8. Cardinal, S., Buchmann, S. L. and Russell, A. L. (2018). The evolution of floral sonication, a pollen foraging behavior used by bees (Anthophila). Evolution 72, 590–600.

9. Clemente, C. J. and Dick, T. J. (2023). How scaling approaches can reveal fundamental principles in physiology and biomechanics. Journal of Experimental Biology 226, jeb245310.

10. Conrad, T. and Ayasse, M. (2015). The role of vibrations in population divergence in the red mason bee, *Osmia bicornis*. Current Biology 25, 2819–2822.

11. Danforth, B. N., Minckley, R. L., Neff, J. L. and Fawcett, F. (2019). The solitary bees: biology, evolution, conservation: Princeton University Press.

12. Darveau, C.-A., Hochachka, P. W., Welch, K. C., Roubik, D. W. and Suarez, R. K. (2005). Allometric scaling of flight energetics in Panamanian orchid bees: a comparative phylogenetic approach. Journal of Experimental Biology 208, 3581–3591.

13. De Luca, P. A., Buchmann, S., Galen, C., Mason, A. C. and Vallejo-Marin, M. (2019). Does body size predict the buzz-pollination frequencies used by bees? Ecology and Evolution 9, 4875–4887.

14. De Luca, P. A., Cox, D. A. and Vallejo-Marin, M. (2014). Comparison of pollination and defensive buzzes in bumblebees indicates species-specific and context-dependent vibrations. Naturwissenschaften 101, 331–8.

15. De Luca, P. A. and Vallejo-Marin, M. (2013). What’s the ’buzz’ about? The ecology and evolutionary significance of buzz-pollination. Current Opinion in Plant Biology 16, 429–35.

16. Delgado, T., Leal, L. C., El Ottra, J. H. L., Brito, V. L. G. and Nogueira, A. (2023). Flower size affects bee species visitation pattern on flowers with poricidal anthers across pollination studies. Flora 299, 152198.

17. Dickinson, M. (2006). Insect flight. Current Biology 16, R309–14.

18. Dickinson, M. H., Lehmann, F. O. and Chan, W. P. (1998). The control of mechanical power in insect flight. American Zoologist 38, 718–728.

19. Dickinson, M. H. and Tu, M. S. (1997). The function of dipteran flight muscle. Comparative Biochemistry and Physiology Part A: Physiology 116, 223–238.

20. Dudley, R. (2000). The Biomechanics of Insect Flight: Form, Function, Evolution: Princeton University Press.

21. Hadfield, J. D. (2010). MCMC Methods for multi-response generalized linear mixed models: The MCMCglmm R Package. Journal of Statistical Software 33, 1–22.

22. Hedtke, S. M., Patiny, S. and Danforth, B. N. (2013). The bee tree of life: a supermatrix approach to apoid phylogeny and biogeography. BMC evolutionary biology 13, 1–13.

23. Hickey, T., Devaux, J., Rajagopal, V., Power, A. and Crossman, D. (2022). Paradoxes of Hymenoptera flight muscles, extreme machines. Biophysical Reviews 14, 403–412.

24. Hrncir, M., Barth, F. G. and Tautz, J. (2005). Vibratory and airborne-sound signals in bee communication (Hymenoptera). In Insect sounds and communication: physiology, behaviour, ecology, and evolution, eds. S. Drosopoulus and M. F. Claridge), pp. 421–436. Boca Raton, Florida: CRC Press.

25. Hrncir, M., Gravel, A. I., Schorkopf, D. L. P., Schmidt, V. M., Zucchi, R. and Barth, F. G. (2008). Thoracic vibrations in stingless bees (*Melipona seminigra*): resonances of the thorax influence vibrations associated with flight but not those associated with sound production. Journal of Experimental Biology 211, 678–685.

26. Jankauski, M., Casey, C., Heveran, C., Busby, M. K. and Buchmann, S. (2022). Carpenter bee thorax vibration and force generation inform pollen release mechanisms during floral buzzing. Scientific Reports 12, 12654.

27. Josephson, R. K. (1997). Power output from a flight muscle of the bumblebee *Bombus terrestris*. III. Power during simulated flight. Journal of Experimental Biology 200, 1241–1246.

28. Kendall, L. K., Rader, R., Gagic, V., Cariveau, D. P., Albrecht, M., Baldock, K. C., Freitas, B. M., Hall, M., Holzschuh, A. and Molina, F. P. (2019). Pollinator size and its consequences: Robust estimates of body size in pollinating insects. Ecology and Evolution 9, 1702–1714.

29. King, M. J. and Buchmann, S. L. (1996). Sonication dispensing of pollen from *Solanum laciniatum* flowers. Functional Ecology 10, 449–456.

30. King, M. J. and Buchmann, S. L. (2003). Floral sonication by bees: Mesosomal vibration by *Bombus* and *Xylocopa*, but not *Apis* (Hymenoptera: Apidae), ejects pollen from poricidal anthers. Journal of the Kansas Entomological Society 76, 295–305.

31. King, M. J., Buchmann, S. L. and Spangler, H. (1996). Activity of asynchronous flight muscle from two bee families during sonication (buzzing). Journal of Experimental Biology 199, 2317–2321.

32. King, M. J. and Lengoc, L. (1993). Vibratory pollen collection dynamics. Transactions of the ASAE 36, 135–140.

33. Kirchner, W. and Röschard, J. (1999). Hissing in bumblebees: an interspecific defence signal. Insectes Sociaux 46, 239–243.

34. Larsen, O. N., Gleffe, G. and Tengo, J. (1986). Vibration and sound communication in solitary bees and wasps. Physiological Entomology 11, 287–296.

35. Ligges, U., Krey, S., Mersmann, O. and Schnackenberg, S. (2018). tuneR: analysis of music and speech. In https://CRAN.R-project.org/package=tuneR.

36. Moore, C. D. and Hassall, C. (2016). A bee or not a bee: an experimental test of acoustic mimicry by hoverflies. Behavioral Ecology 27, 1767–1774.

37. Pritchard, D. J. and Vallejo-Marin, M. (2020). Floral vibrations by buzz-pollinating bees achieve higher frequency, velocity and acceleration than flight and defence vibrations. Journal of Experimental Biology 223, jeb.220541.

38. Russell, A. L., Buchmann, S. L. and Papaj, D. R. (2017). How a generalist bee achieves high efficiency of pollen collection on diverse floral resources. Behavioral Ecology 28, 991–1003.

39. Snodgrass, R. E. (1935). Principles of Insect Morphology. New York: McGraw-Hill.

40. Snodgrass, R. E. (1942). The skeleto-muscular mechanisms of the honey bee. Smithsonian Miscellaneous Collections 103, 1–120.

41. Sueur, J. (2018). Sound Analysis and Synthesis with R. The Netherlands: Springer.

42. Team, R. D. C. (2023). R. A language and environment for statistical computing. R Foundation for Statistical Computing, Vienna, Austria. URL http://www.R-project.org, vol. 4.3.1.

43. Vallejo-Marin, M. (2022). How and why do bees buzz? Implications for buzz pollination. Journal of experimental botany 73, 1080–1092.

44. Vallejo-Marín, M. and Vallejo, G. C. (2020). Comparison of defence buzzes in hoverflies and buzz-pollinating bees. Journal of Zoology 313, 237–249.

